# Testis-expressed cluster of microRNAs 959-964 controls spermatid differentiation in *Drosophila*

**DOI:** 10.1101/013243

**Authors:** Sergei Ryazansky, Elena Mikhaleva, Natalia Akulenko, Oxana Olenkina

**Affiliations:** Institute of Molecular Genetics, Kurchatov sq. 2, 123182 Moscow, Russia

**Keywords:** microRNA, miR-959-964, spermatogenesis, testes, drosophila

## Abstract

MicroRNAs are a wide class of ∼22 nt non-coding RNAs of metazoans capable of inhibiting target mRNAs translation by binding to partially complementary sites in their 3’UTRs. Due to their regulatory potential, miRNAs are implicated in functioning of a broad range of biological pathways and processes. Here we investigate the functions of the miR-959-964 cluster expressed predominantly in testes of *Drosophila melanogaster*. The deletion of miR-959-964 resulted in male sterility due to the disturbance of the spermatid individualization process. Analysis of the transcriptome by microarray followed by luciferase reporter assay revealed *didum, for, fdl* and CG10512 as the targets of miR-959-964. Moreover, the deletion of miR-959-964 is accompanied by a decreased the expression of genes responsible for microtubule-based movement and spermatid differentiation. Thus, we suggest that miR-959-964 can control the process of spermatid individualization by direct and indirect modulating the expression of different components of the individualization process. In addition, we have shown that in comparison to other miRNAs, the rate of evolution of the testis-specific miR-959-964 cluster is unusually high, indicating its possible involvement in speciation via reproductive isolation.

## Introduction

MicroRNAs (miRNAs) are short 21-23 nt RNAs that are processed by Dicer and Drosha proteins from 5’- and 3’-end arms of 60-100 nt precursors having a characteristic hairpin-like secondary structure (pre-miRNAs). The genes of miRNAs are often located close to each other in the genome forming polycystronic clusters [1–3]. The most amazing feature of miRNAs is their ability to repress the expression of target genes on the post-transcriptional level. In animals, the recognition of the 3’ untranslated regions (3’UTRs) of targets by partially complementary miRNAs causes mRNA degradation and/or blocking of translation, which is mediated by Argonaute (Ago) and other effector proteins [4–7]. The key region of a miRNA that is responsible for the recognition of target mRNAs is the so called ‘seed’ region embracing nucleotides 2 to 7 relative to its 5’-end (Bartel, 2009). In geneal, miRNA-mediated inhibition of target mRNAs is a powerful and flexible mechanism of tissue-specific regulation of gene expression and, as a result, the different pathways and processes.

The involvement of miRNAs in development and functioning of the male reproductive system is shown in a number of reports (see Reviews [8, 9] for futher details). The first indication of this possibility was the demonstration that Dicer, Ago and several testis-expressed miRNAs are co-localized in the chromatoid bodies of male germinal cells [10]. The deletion of *dicer* or *drosha* associated with the decline of miRNA abundance results in the failure of spermatogenesis in mammals [11–15]. It is known that mammalian testes have a specific set of miRNAs [16–21]. Murine testis-specific miRNAs are mostly expressed in pachytene spermatocytes [19], and indeed the ablation of Dcr1 specifically impaired the meiotic and post-meiotic stages of spermatogenesis which finally led to infertility [14]. In some reports specific functions of particular miRNAs during spermatogenesis were demonstrated. miRNAs from the miR-17-92 cluster are able to inhibit the translation of E2F1 mRNA resulting in prevention of apoptosis in meiotic spermatocytes [22]. miRNA-21 regulates the self-renewal of germ stem cells of murine testes [23]. In somatic cells of the niche in testes of old-aged fruit flies let-7 represses the expression of *Imp* that is implicated in maintaining germ stem cells divisions [24]. It has recently been demonstrated that miR-34 and miR-449 repress E2F, NOTCH1 and BCL2 in mammalian testes [20, 25, 26]. In addition to this, miR-34c is able to enforce apoptosis of germ cells [27], and, being transmitted to the oocyte by sperm, is implicated in the initiation of the first zygotic division [28]. Murine *Tnp2*, encoding a transition protein involved in chromatin remodeling during spermatogenesis, is regulated by miR-122a [21].

It is known that mid spermatogenesis is characterized by a complete cease of gene transcription, and therefore proteins required for later stages of spermatid differentiation must be produced by translation of mRNAs stored prior to the cease of transcription [29, 30]. Translation of such mRNAs is blocked at the mid stage and activated at the late stages, while their premature translation activation results in abnormal spermatogenesis and infertility. The described mechanisms of translational repression that can occur in meiotic and post-meiotic cells include the binding of repressor proteins to the Y-box, poly(A) signals or AU motifs in the 3’UTR of mRNAs, and also the loading of mRNAs into ribonucleic protein particles (RNPs) that prevents them from degradation [29, 30]. Although the instances of miRNA functioning in early spermatogenesis are known, the degree of the contribution of miRNAs to post-transcriptional regulation of gene expression during late spermatogenesis is still under the question. Here we present the data demonstrating the possibility of the involvement of a whole cluster of miRNAs in the regulation of spermatid differentiation in testes of *Drosophila melanogaster*.

## Materials and Methods

### Analysis of small RNA libraries

Tissue expression profiling of miRNAs was conducted with small RNA libraries fetched from GEO: GSM239041 (heads), GSM278695 (males), GSM278706 (females), GSM280085 (testes), GSM280082 (ovaries), GSM286604 (0-1h embryos), GSM286605 (2-6h embryos), GSM286607 (6-10h embryos) and GSM272652 (S2). Small RNA reads have been mapped to dm3 genome assembly using *bowtie* [31] with the requirement of perfect matching. Reads were annotated according to UCSC Genome Browser [32] and miRBase [33] databases, and the counts of miRNAs were used to infer their expression profiles.

### Northern-blot hybridization

A total of 20 μg of RNA extracted from fly heads, germinal tissues, carcasses (bodies without germinal tissues), larvae, embryos and S2 cells, were resolved on 20% PAGE in denaturing conditions, transferred to Hybond N+ membrane (Amersham-Pharmacia Biotech, GE Healthcare Bio-Sciences) and fixed by UV radiation (Stratagene). Hybridization of 10 pmol of [γ-^32^ P]ATP labeled probes (Table S1) was performed in Church buffer overnight at 37°C with subsequent double washing in 2x SSC/0.1% SDS. The signals were visualized with the STORM PhosphorImager System (Amersham-Pharmacia Biotech, GE Healthcare Bio-Sciences).

### FRT-FLP mediated deletion

The deletion of the region encompassing the *miR-959-964* cluster was performed as described previously [34]. Fly stocks *f07682* and *f02063*, carrying *PiggyBac*-based transgenic constructs with FRT sites and flanking the region of interest, were obtained from Harvard Stock Center. Fly stock *#279* from Bloomington Stock Center was used as the source of FLP recombinase. The expression of FLP was activated by heat-shock treatment of flies (1h, 37°C) for five days. F2 progeny was screened for the presence of the deletion by site-specific PCR, as described [34] (primers as in Table S1). The genotype of the obtained fly strain after isogenisation was *iso w^1118^; ΔmiR-959-964*/*CyO; iso3*. Flies with *iso w^1118^; ΔmiR-959-964/CyO; iso3* and *iso w^1118^; ΔmiR-959-964/ΔmiR-959-964; iso3* genotypes are designated throughout the text as ‘wild type’ and *ΔmiR-959-964*, respectively.

### Fertility test

15 3-day old males of the tested genotype were individually crossed with five 3-day old virgin females of the control *Df(1)yw^67c23(2)^* strain. The parents were removed from vials after 5 days of crossing. The level of male fertility refers to the average number of hatched progeny obtained from the each cross.

### Fly stocks

The transgenic fly strains ID#102675 and ID#100781 encoding the shRNA hairpins targeted CG18266 and CG31646 respectively were obtained from VDRC [35]. The fly strain expressing *ProtB-eGFP* was kindly provided by R. Renkawitz-Pohl and C. Rathke [36]. For testing rescue transgenic constructs the following Gal4 sources were used: ubiquitously-expressed *tubP-GAL4* (Bloomington Stock No. 5138); early-germline expressed *nos-GAL4* (Bloomington Stocks No. 7303, 27571 and 31777); testis-soma expressed *tj-GAL4* (Bloomington Stock No. 50105 and 50165) and *C587-GAL4* (kindly provided by M. Buszczak); *P{w[+mGT]=GT1}BG01278* (Bloomington Stock No. 12608) and *P{w[+mGT]=GT1}brat[BG02721]* (Bloomington Stock #12820) active in late spermatogonia and early spermatocytes. The other fly strains are: Mi{ET1}CG31646^MB09592^ (Bloomington Stock No. 27790), Mi{MIC}CG31646^MI03191^ (Bloomington Stock No. 35926). Fly strains were grown under standard conditions at 25°C with the only exception of *C587-GAL4* strain that was grown on 18°C.

### Immunofluorescence and phase-contrast microscopy

Immunostaining of testes was performed as described [37] using mouse anti-histone (1:1200, Millipore) as primary antibodies and anti-mouse Alexa-Fluor-647 (Invitrogen) as secondary antibodies. F-actin was stained by rhodamine phalloidin conjugate. Immunofluorescence and eGFP samples were examined using laser scanning confocal LSM 510 META microscope (Carl Zeiss) equipped with appropriate filters. Phase-contrast microscopy of slides with squashed testes was performed as described previously [38] on DM6000B microscope (Leica).

### Microarray

Total RNA from testes of 3-5 day old males extracted using TRIzol reagent (Gibco) was treated with DNase I (Ambion) and reverse transcribed with SuperScript III (Invitrogen). After synthesis of the second strand using SuperScript Choice System (Invitrogen), double-stranded cDNA was labeled with Cy3 or Cy5 dyes by using Dual-Color DNA Labeling Kit (NimbleGen, Roche Applied Science) and purified on microcon-30kDa (Amicon). Equal amounts (5-10 μg) of Cy3-cDNA from wild type and Cy5-cDNA from *ΔmiR-959-964* testes were mixed, dried and re-dissolved in 10 μl of DIG Hyb buffer (Roche Applied Science) containing 100 μg of yeast tRNA (Invitrogen) and 100 μg of salmon sperm DNA (Ambion). Hybridization was performed overnight at 37°C on FL003 microarrays (FlyChip) in the Array Hybridization Cassette (Invitrogen) in a water bath.

Immediately after slide washing by the Array Wash Buffers (NimbleGen), they were scanned using the GenePix 4400B microarray scanner (Molecular Devices). Each array was repeated twice with dye swap. The images with intensity signals were evaluated in GenePix Pro 6.0 (Molecular Devices) followed by analysis using *limma* package of R/Bioconductor [39]. Only spots with signal-to-noise ratio ≥3 in both channels were taken into account. After subtraction of background from signals, they were normalized within each array and between arrays by using *loess* and *quantile* methods, respectively. Microarray data are deposited in the ArrayExpress repository (E-MTAB-3214). oligoMath program from Kent utilities was used for the search of motifs in the 3’UTRs that were complimentary to the ‘seed’ regions of miRNAs. The analysis of Gene Ontology terms and KEGG pathways enrichment was evaluated using FlyMine project, v37.0 [40].

### RT-qPCR

Total RNA from testes of 3-5 day old males extracted using TRIzol reagent (Gibco) was treated with DNase I (Ambion) and reverse transcribed with SuperScript II (Invitrogen). Quantitative real-time PCR of cDNA was conducted on DT-96 (DNA Technology, Russia). Each PCR was performed with two technical replicates, while final mean and standard deviations values were calculated on three biological replicates. The expression of house-keeping *Rp49* gene was used for normalization.

### Dual-luciferase assay

Cluster *miR-959-964*, fusions of *Renillia* luciferase *RLuc* with 3’UTR fragments of genes containing the miRNA recognitions sites, and firefly luciferase *Luc* were cloned under the control of *actin* promoter into pAc5.1/V5-His B vector (Invitrogen). Primers used for the amplification of *miR-959-964* and 3’UTR fragments from genomic DNA are listed in Table S1. Three obtained plasmids encoding *miR-959-964*, *RLuc*-3’UTR and *Luc* were co-transfected using X-tremeGENE DNA reagent (Roche Applied Science) into S2 cell culture in 100:25:25 ng per well amounts, respectively. Each treatment was performed in triplicate in 96-well plates. The measurement of luminescence was performed after 48h of transfection with Dual-Luciferase Reporter Assay System (Promega) on Modulus Microplate Mulimode Reader (Turner BioSystems). Each assay was repeated three times.

### Cloning of miR-959-964 rescue transgenic constructs

We have constructed two transgenic vectors contained either the whole *miR-959-964* cluster (∼1.6 kb in total, pUAST-miR959-964), or pre-miR-963, pre-miR-964 and ∼4 kb upstream region (∼4.3 kb in total, pUAST-pr.miR963-964). The genomic DNA was amplified by clu_d2/clu_r5 (pUAST-miR959-964), or CG31646_1/CG31646_2 and CG31646_3/CG31646_4 pairs of primers (pUAST-pr.miR963-964, sequences of primers in Table S1) by using high-fidelity polymerase (Evrogen). PCR amplicons were extracted, purified and digested by *Xho*I and *Xba*I (pUAST-miR959-964), and *EcoR*I and *Not*I or *Not*I and *Xba*I (pUAST-pr.miR963-964). The cloning of PCR fragments was performed into the pUASTattB vector that is designed specifically for using in the phiC31 Drosophila transgenesis system [41]. For pUAST-pr.miR963-964 firstly CG31646_1/CG31646_2 fragment was cloned followed by cloning of CG31646_3/CG31646_4 fragment.

### Obtaining of transgenic fly strains

Transgenic fly strains carrying either pUAST-miR959-964 or pUAST-pr.miR963-964 transgenes were obtained by using phiC31 site-specific integration system [41]. For transgenesis strains with attP landing sites *68E1* (Bloomington Stock #24485; pUAST-miR959-964) or *75A10* (Bloomington Stock #24862; pUAST-miR959-964, pUAST-pr.miR963-964) were used. After balancing, the transgenic strains were used for obtaining the fly strains carrying rescue transgene (pUAST-miR959-964 or pUAST-pr.miR963-964), *ΔmiR959-964* deletion and transgene expressing one of the tissue-specific Gal4 drivers (see Result section for details). The males (10-50) of these strains were used for fertility test by crossings with virgin 3-5 day-old *Df(1)yw^67c23(2)^* females. In strains carrying *tubP-GAL4* and *P{w[+mGT]=GT1}BG01278* drivers we did not receive any adult males while for *C587-GAL4* driver only few males were obtained.

### Phylogenetic reconstruction of cluster evolution

Multiple alignments of pre-miRNA encoding sequences from *Drosophilidae* species were extracted from UCSC Table Browser [32, 42]. After manual evaluation of correctness of each alignment, they were merged into one cluster alignment which was submitted to the PHYLIP program [43] for estimating the distances between species and construction of the phylogenetic tree. The stabilities of secondary structures of pre-miRNAs were evaluated by using RNAfold [44] followed by randfold [45]. The rate of nucleotide substitution was measured by the Kimura model [46] implemented in the BioPerl Toolkit [47].

## Results

### miR-959-964 cluster is expressed in testes

To identify the miRNAs that are presumably expressed in fly testes we reanalyzed the set of publicly available small RNA libraries from heads, bodies, testes, ovaries, embryos and S2 cells of *D. melanogaster* (as indicated in the Method section). Applying the strong requirement of at least 4-fold enrichment of the abundance of miRNAs in the library from testes relative to any other analyzed tissue, allowed us to reveal 22 miRNAs expressed predominately in testes (Fig. 1A). Testis-specific miRNAs include miRNA-316, miRNA-375, miRNA-982, miRNA-983, miRNA-985, miRNA-1004 and also 16 miRNAs from miR-959-964, miR-972-979 and miR-991-992 clusters. Almost all of these miRNAs are found only in insects. The only exception is miRNA-375 that is conserved among animals, and its expression in testes seems to be specific for flies since in mammals it is known as a tumor suppressor expressed in a broad range of tissues [48].

**Figure 1.**
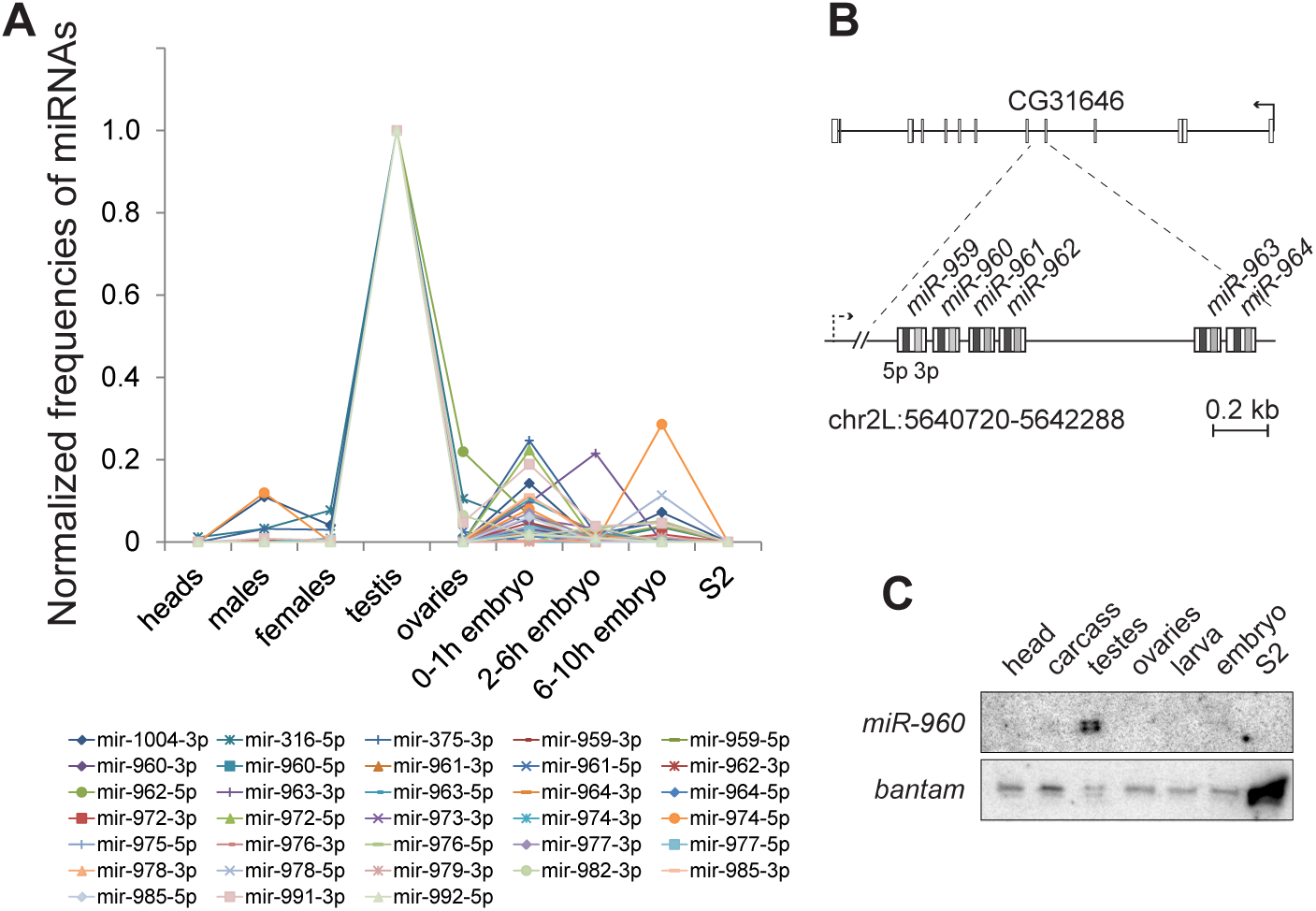
Testis-specific miRNAs of *Drosophila*. (A) Tissue expression profiles of testis-specific miRNAs of *Drosophila*. The frequencies of each miRNA in each tissue were normalized to its frequency in the testes. (B) The structure of the *miR-959-964* cluster in *D. melanogaster*. Broken dashed arrow indicates the proposed TSS of cluster pri-pre-miRNA. (C) Tissue expression profile of *dme-miRNA-960* revealed by Northern-blot analysis. The hybridization on ubiquitously-expressed *miRNA-bantam* was used as loading control.

The testis-specific character of the miRNAs’ expression implies their possible important roles during spermatogenesis. To investigate the functions of miRNA clusters during spermatogenesis, we chose to focus specifically on the miR-959-964 cluster. This cluster is encoded on the opposite DNA strand of the intronic region of the protein-coding gene CG31646, spanning ∼1.2 kb and encompassing six miRNAs, including miRNA-959, miRNA-960, miRNA-961, miRNA-962, miRNA-963, and miRNA-964 (Fig. 1B). The transcription of the cluster is probably initiated at a putative transcription start site located at a distance of ∼4 kb upstream (Table S3 of [2]). Although it has recently been reported that miR-959-964 is expressed at some level in *Drosophila* heads where it is involved in the circadian control of feeding [49], our profiling of its tissue expression by Northern-blot (Fig. 1C) as well by the analysis of small RNA libraries (Fig. 1A; Fig. S1) clearly showed that miR-959-964 cluster is expressed predominantly in the testes of adult flies.

### Deletion of *miR-959-964* causes male sterility and a failure in spermatid individualization

To elucidate possible functions of miR-959-964 during spermatogenesis we constructed a mutant fly strain carrying the deletion of the cluster. The deletion was produced by FLP-directed recombination between flanking *PiggyBac*-based transgenic constructs carrying *FRT* sites (Fig. S2A) [34]. To test the proposition that the absence of testis-specific miR-959-964 can affect spermatogenesis, we inspected the level of male fertility. Indeed, we have found that males, but not females, carrying this deletion are completely sterile (Fig. S2B, data not shown). The deletion also encompassed three other adjacent protein-coding genes with unknown functions (Fig. S2A). According to the modENCODE data [50], CG14010 and CG31646 not expressed in adult testes at a detectable level (data not shown). Moreover, we have performed the *nanos*-driven RNAi-knockdown of the CG18266 and CG31646 and found that it didn’t influence male fertility (Fig. S2C). We have also tested the male fertility of the couple of mutant fly strains carrying insertions of transgenes upstream or within miR-959-964 cluster. We found that males of mutant Mi{ET1}CG31646^MB09592^ carrying the insertion of a transgene in CG31646 and upstream of miR-959-964 are fertile (Fig. S2C) while 0-2 days old males carrying the Mi{MIC}CG31646^MI03191^ insertion within miR-959-964 cluster are sterile (Fig. S2D). Thus, male sterility seems to be caused by the deletion of *miR-959-964* but not any of the adjacent protein-coding genes.

In *Drosophila* testes the progeny of the first division of germinal stem cells (spermatogonium) is encapsulated by somatic support cells forming a cyst. The spermatogonium proceeds through the series of mitotic and meiotic divisions resulting in 64 round spermatids. During the following differentiation, the spermatids are tailed, and the histones of elongating nuclei are replaced by transition proteins, which are subsequently replaced by protamine-like proteins. And finally, since the mitotic and meiotic divisions are characterized by incomplete cytokinesis, the elongated spermatids within the encysted bundles undergo the process of individualization [51]. To determine the nature of sterility of *ΔmiR-959-964* males, we inspected mitotic divisions, meiotic divisions and spermatid differentiation stages of spermatogenesis using phase-contrast microscopy of squashed testes. We didn’t observe any noticeable alterations in mitosis and meiosis in testes of *ΔmiR-959-964* flies, and they contain all cell types that can be expected after successful proceeding of divisions (Fig. S3). At the same time, we found that deletion of *miR-959-964* is accompanied by a failure of the last stage of spermatogenesis i.e. of the process of spermatid differentiation. In wild type testes the encysted spermatids are grouped in orderly and regularly structured bundles, enveloped by shared somatic support cells of the cyst. In contrast, in the case of the *miR-959-964* deletion the spermatid bundles became looser with a clear tendency to lose the ordered structure of their organization (Fig. 2A; Fig. S3). Additionally, although the nuclei of the wild type spermatids are bundled within cysts at distal ends of testes (Fig. S4A,B), the nuclei of *ΔmiR-959-964* spermatids appear to have the so called ‘scattered’ phenotype (Fig. S4C,D,E) that is typical for mutants with abnormal spermatid differentiation [52].

**Figure 2.**
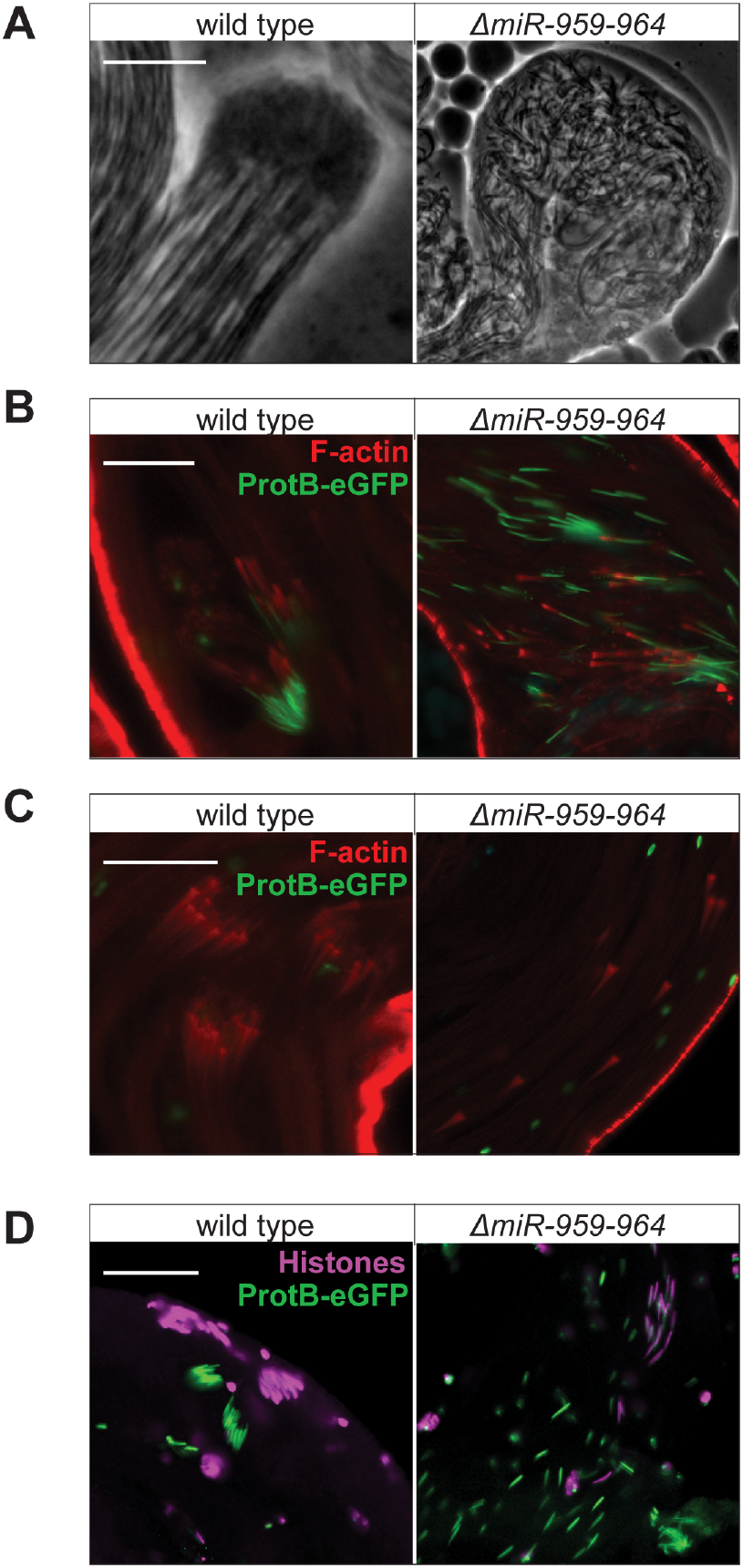
Spermatid individualization in wild type and *ΔmiR-959-964* testes. (A) Phase-contrast microphotographs of nuclei-contained heads of spermatid bundles. Scale bar, 10 μm. (B,C) Confocal microscopy of the phalloidin-stained investment cones (ICs, red) of spermatids near their nuclei (ProtB-eGFP, green) (B) and on the tails (C). (C) represents more later stage of the individualization process in comparison to (B). The wild type ICs are come together in compact bundles and synchronically moved along the tails; the ICs in *ΔmiR-959-964* testes didn’t congregate in compact bundles and moved along the tails independently of each other. (D) Confocal microscopy of ProtB-eGFP and immunostained histones in the nuclei of elongated spermatids. (B,C,D) Scale bar, 20 μm.

The individualization process of the encysted and elongated spermatids includes the involvement of the actin-rich investment cones (ICs), assembled near each nucleus. During individualization, these structures are congregated with each other in a tight individualization complex and moved downward the spermatid axonema in a coordinated manner, accompanied by the formation of the separated individual sperm cells membranes and extruding the excess of the cytoplasm into the terminal waste bag. The loss of ordered organization of *ΔmiR-959-964* spermatid bundles may indicates that their individualization has not occurred properly. To test this possibility, we performed the staining of the ICs of the whole-mount testes by phalloidin. Indeed, we have found that although the ICs are appeared to assemble normally in testes of *ΔmiR-959-964* flies, they are not come together and moved asynchronically and independently of each other (Fig. 2B,C).

We also inspected the initial stage of the spermatid individualization in *ΔmiR-959-964* testes. Normally, the process of spermatid individualization onsets after the replacement of histones to the protamine-like proteins in the nuclei of elongated spermatids, which is associated with the chromatin condensation and changing of the nucleus shape, from an elliptic to a needle-like one. By using the *ProtB-eGFP* transgenic construct encoding the fusion of protamine B and eGFP [53] and immunostaining of histones, we have shown that although in the testes of *ΔmiR-959-964* flies the replacement of histones occurred similar to wild type flies, spermatids either with histone- or protamine- containing nuclei lost their ordered packaging in the bundle-like structures (Fig. 2D). This finding can be explained by premature launching of the individualization process of mutant spermatids before the replacement of histones to protamine-like proteins. The alternative explanation is that the observed failure of coordinated individualization is the result of the loss of correct alignment of germ cells within cyst right after meiotic divisions. Since we didn’t find any strong evidences that ICs are able to be formed prior to the histones to protamine replacement in elongated spermatids of *ΔmiR-959-964* testes (data not shown), we compelled that the second proposition is more plausible.

### Identification and characterization of miR-959-964 targets

Having identified significant changes in the phenotype of the mutant strain, we sought to determine the putative targets of the miR-959-964 cluster. Assuming that deletion of miRNAs will cause the target up-regulation, we compared the transcriptomes of testes from wild type and *ΔmiR-959-964* flies using the microarray technique. We have shown that the deletion of *miR-959-964* resulted in the up-regulation of 15 and down-regulation of 43 genes in testes (at least 2-fold change, *Padj-value*≤0.05) (Table 1; Table S2). To determine the possible targets of miR-959-964, we searched sites complementary to the ‘seed’ region of miRNA (2-7 nt) in the 3’UTRs of up-regulated genes. Indeed, we succeeded in finding that more than half of the up-regulated genes (9 among 15) contain putative recognition sites for −3p and −5p miRNAs (Table 1). The number of sites per 3’UTR varied from a single one to six for up to four different miRNAs (Table 1). The gene *didim* has the largest number of miRNA recognition sites in its 3’UTR (6 sites for miRNA-959-3p, miRNA-962-5p, miRNA-963-5p, and miRNA-964-5p) and it is also predicted as miR-959-964 target by TargetScanFly 6.2 program [54]. Other up-regulated genes include CG1597 with three recognition sites of miR-959-964, *fdl*, *for,* CG10512 and CG8924 with two sites, and *debcl*, *Cyp12d1-d*, CG2617 with one site (Table 1).

**Table 1.** Protein-coding genes with increased level of expression in testes of *ΔmiR-959-964* flies (revealed by microarray). The adjustment of *P-values* for multiple hypothesis correction was evaluated with Benjamin-Hochberg algorithm. The −5p and −3p miRNAs of miR-959-964 that have putative recognition sites in the 3’-UTRs of corresponding genes are also listed.

| Gene | logFC | *P-value* | *P<sub>adj</sub>-value* | miRNAs |
| --- | --- | --- | --- | --- |
| CG30053 | 2.11 | 1.48E-09 | 1.16E-06 |  |
| CG18477 | 1.70 | 1.02E-06 | 1.33E-04 |  |
| CG8924 | 1.62 | 3.12E-06 | 3.05E-04 | miRNA-959-5p, miRNA-963-3p |
| *Cyp12d1-d* | 1.42 | 4.41E-05 | 2.66E-03 | miRNA-963-3p |
| CG1597 | 1.29 | 2.03E-04 | 9.75E-03 | miRNA-960-5p, miRNA-961-5p, miRNA-964-5p |
| *fdl* | 1.28 | 2.28E-04 | 1.07E-02 | miRNA-959-3p, miRNA-962-5p |
| *debcl* | 1.28 | 2.34E-04 | 1.08E-02 | miRNA-964-5p |
| *didum* | 1.27 | 2.11E-07 | 4.12E-05 | miRNA-959-3p, miRNA-962-5p (2 sites), miRNA-963-5p, miRNA-964-5p (2 sites) |
| CG3781 | 1.20 | 5.42E-04 | 2.16E-02 |  |
| CG2617 | 1.13 | 4.08E-06 | 3.55E-04 | miRNA-964-3p |
| CG10512 | 1.12 | 1.19E-03 | 3.66E-02 | miRNA-960-5p, miRNA-964-3p |
| CG10887 | 1.11 | 1.34E-03 | 3.94E-02 |  |
| *for* | 1.11 | 1.41E-03 | 4.00E-02 | miRNA-964-3p, miRNA-964-5p |
| CG18598 | 1.10 | 1.51E-03 | 4.23E-02 |  |
| CG8549 | 1.05 | 1.80E-05 | 1.36E-03 |  |

To further confirm the up-regulated genes as the genuine target genes, we performed the dual-luciferase assay. For this, the fragments of the 3’UTRs containing the miRNA recognition sites of the corresponding genes were cloned downstream of *Renillia* luciferase *RLuc* gene driven by the strong constitutive the actin promoter and co-transfected with the *miR-959-964* cluster under the *actin* promoter in the S2 cell line. For normalization of the luminescence between replicas we used a vector expressing the Firefly luciferese *Luc*, that was also co-transfected in S2 cells. As a negative control either the vector expressing the anti-sense strand of the miR-959-964 cluster or the empty vector without the cluster were used. Using this system, we succeeded to confirm that the expression of tested constructs containing 3’UTRs of *for*, *didum*, CG10512 and *fdl* is repressed in the presence of miRNA encoding plasmid (Fig. 3A). The up-regulation of candidate targets in *ΔmiR-959-964* testis is confirmed by RT-qPCR (Fig. 3B) and therefore indeed they can be considered as the miR-959-964 targets.

**Figure 3.**
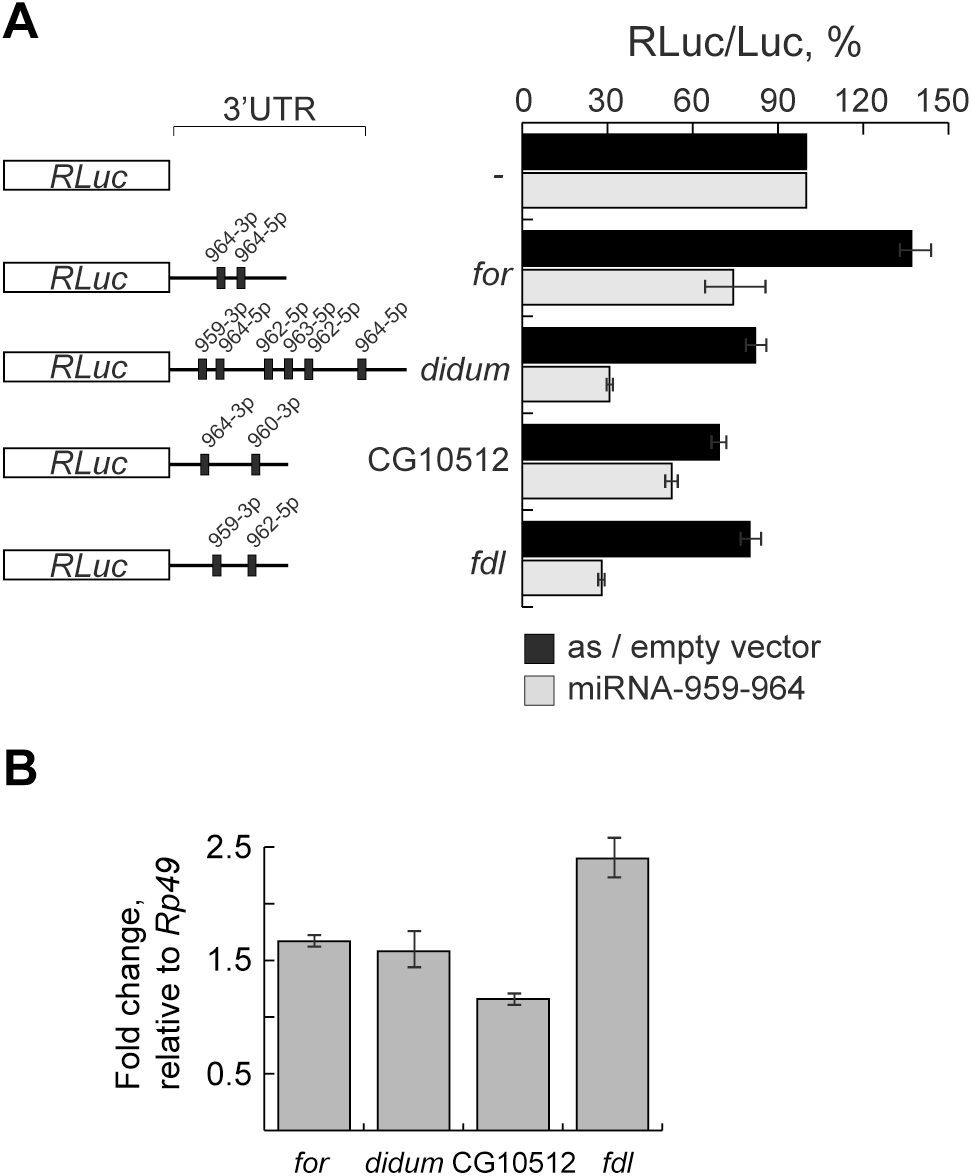
The regulation expression of miR-959-964 targets. (A) The dual-luciferase assay for testing of miR-959-964 targets. The S2 cells were co-transfected by three vectors expressed *RLuc-3’UTR*, *Luc*, and either *miR-959-964* (black bars) or control vector (gray bars). Control vector was either without any CDS (‘empty’) or carried the miR-959-964 sequence in the anti-sense orientation. (B) Fold change of steady-state level of expression of the indicated gene targets in *ΔmiR-959-964* testes relative to wild-type ones. The fold change of *for*, *didum* and *fdl* genes expressions are statistically significant (*P-value* ≤ 0.05).

Additionally, to identify which pathways had a potential to be most influenced by the *ΔmiR-959-964* deletion, we performed the analysis of enrichment of Gene Ontology terms and KEGG pathways among the up- and down-regulated genes revealed by microarray. Since the functions of most up-regulated genes are not described, we succeeded in finding the enrichment among GO terms only for down-regulated genes (Table S2). Specifically, the biological functions of most of them are related to microtubule-based movement and the spermatid differentiation processes, and also they are annotated as the components of the axonemal dynein complex, which is responsible for the assembly and movement of the ICs (Table S3).

### Germline and somatic expression of *miR-959-964* transgene on early-stages of spermatogenesis dose not rescue the sterility of *ΔmiR-959-964*

In the order to check on which stage of spermatogenesis and in what tissue miR-959-964 can be active, we obtained the transgeneic fly strains carrying the transgene with the whole miRNA-959-964 under the control of UAS element. To test if this construct can rescue *ΔmiR-959-964* phenotype, its expression was activated in testis of *ΔmiR-959-964* males by using the different sources of GAL4 driver. Such, we tested ubiquitously-expressed *tubP-GAL4,* spermatogoium - expressed *nos-Gal4*, late-spermatogonium and early-speramtocyte expressed *P{w[+mGT]=GT1}BG01278* and *{w[+mGT]=GT1}brat[BG02721]*, early-soma expressed *tj-Gal4* and *C587-Gal4* drivers. The expression of *tub-GAL4* results to fly lethality on the very early stages of embryogenesis, that is consistent with the early observation where hypexpression of miR-959-964 leads to the fly lethality [55]. In all of other cases we didn’t observe the rescue of *ΔmiR-959-964* males sterility by transgene.

Recently it was shown that deletion of only two miRNAs from cluster – miRNA-963 and −964 – results to male sterility [56]. To test, if the expression of only two of these miRNA can rescue the mutant *ΔmiR-959-964* phenotype, we have constructed the second transgenic construct encoding miRNA-963 and −964 and containing long ∼4 kb upstream region encompassing the predicted promoter of miR-959-964. Similarly to the transgene encoding the whole miR-959-964 cluster, in this case we didn’t observe the rescue of fertility of *ΔmiR-959-964* males by the using the above set of drivers. This construct alone, without any GAL4 driver, also did not rescue the mutant phenotype indicating that besides upstream 4kb region other *cis*-regions may be necessary for miR-959-964 expression in male testis. In summary, these results show that expression of miR-959-964 cluster may occur on the late stages of spermatogenesis, where none of the above used GAL4 drivers are expressed.

### *miR-959-964* undergoes rapid evolution

The miR-959-964 cluster is unique to the *Drosophilidae* genus and absent in any other animals. All miRNAs from miR-959-964 can be attributed to different families since they are not related to one another or to any other drosophila miRNAs and also have different ‘seed’ regions (Fig. 4A). This observation suggests that formation of *miR-959-964* has occurred without duplication of pre-miRNAs emerged within the cluster independently of each other.

**Figure 4.**
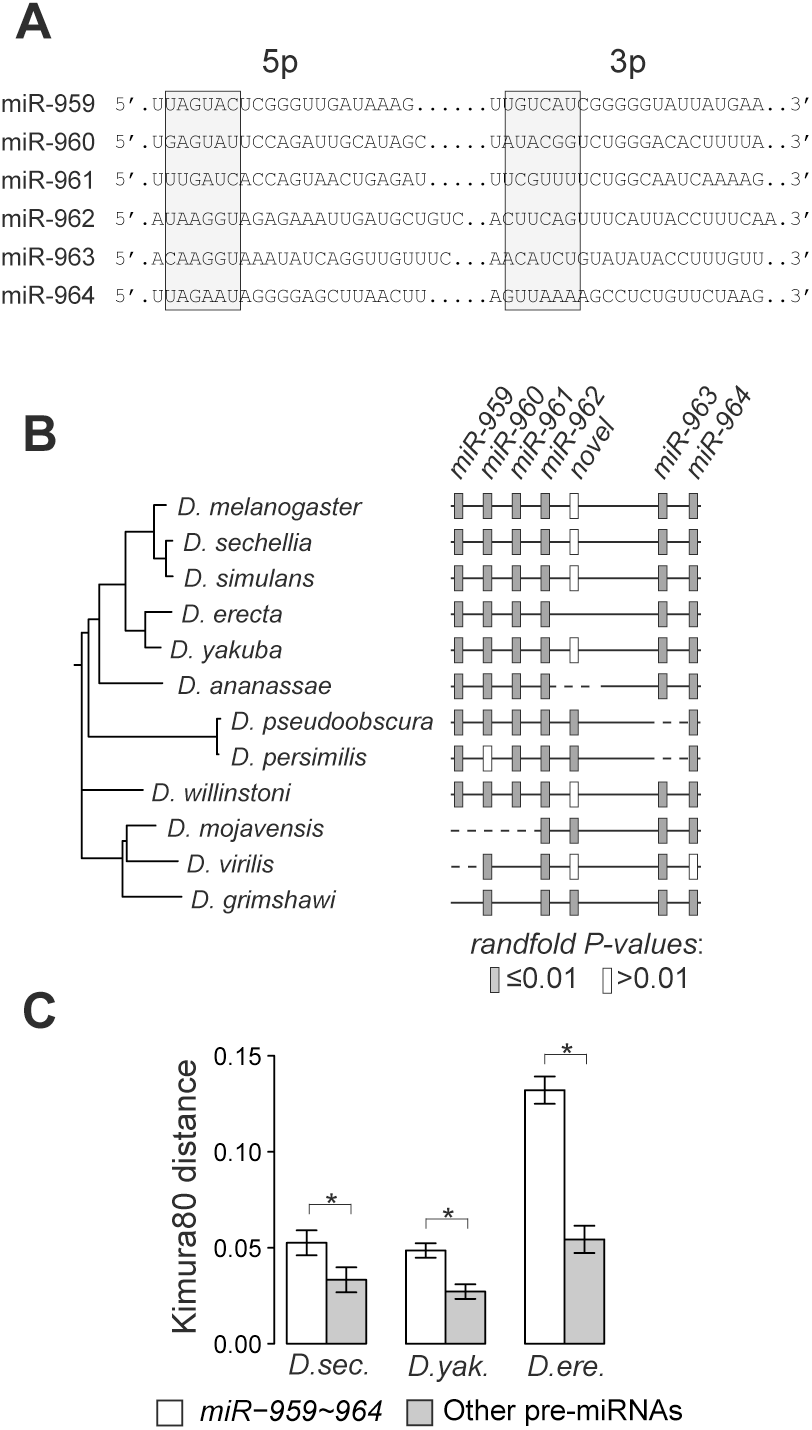
The phylogeny of *miR-959-964.* (A) The sequences of −5p and −3p miRNAs of the miR-959-964. The ‘seed’ regions are highlighted by gray boxes. (B) The phylogenetic reconstruction of *miR-959-964* structure in *Drosophilidae* genus. *P-value* evaluated by randfold [45] reflects the probability that minimum free energy of the given secondary structure is different from one of the random structures, and only hairpins with significant *P-values* can be considered as energetically stable ones (gray color). (C) The rate of nucleotide substitutions in pre-miRNAs of *miR-959-964* relative to other *Drosophila* pre-miRNAs. The measurement of substitution rate was performed by applying the Kimura substitution model [43] to the corresponding pre-miRNA sequences from pair of species, *D. melanogaster* vs. *D. sechellia*, *D. yakuba* or *D. erectus*, as indicated in the plot legend. Error whiskers represent standard errors; significant differences between rate means was evaluated by one-sided Student’s t-test (*P-values* are **<0.01 and *<0.05).

It has been reported earlier that mammalian testis-expressed miRNAs similarly to protein-coding genes are characterized by a heightened rate of nucleotide substitutions that reflect their rapid evolution [57, 58]. In order to extend this observation for fly miRNAs, we reconstructed the phylogenetic history of the whole *miR-959-964* cluster. We found that the reconstructed tree of the cluster is highly similar to the phylogenetic tree of the *Drosophilidae* genus indicating the coincidence of *miR-959-964* evolution with the evolution of the fruit flies (Fig. 4B). At the same time, although the structure of the *miR-959-964* in *Drosophilidae* is almost the same within the genus, it have undergone notable changes during evolution (Fig. 4B,C). First, the size of *miR-959-964* varies from 4 to 7 miRNAs in the different species. Second, in some of the species the clustered pre-miRNAs lost their ability to fold into the stable hairpin-like secondary structures due to mutational process (Fig. 4B). For example, in *D. pseudoobscura* and several other species we identified pre-miRNA-like sequence encoded between *miRNA-962* and *miRNA-963* (Fig. 4B; Fig. S5). In *D. pseudoobscura* this pre-miRNA-like sequence has a valid and stable secondary structure indicating its opportunity to be processed into the functional mature miRNAs. At the same time, the ortholog sequence of *D. melanogaster* is predicted to have no stable hairpin secondary structure (Fig. S5) that explains its missing in the previous genome-wide miRNA identification screens. Finally, we measured the rates of sequence evolution, and found that pre-miRNAs of the *miR-959-964* cluster in contrast to other pre-miRNAs have significantly higher rate of nucleotide substitutions (Fig. 4C). Taken together, these observations show that *miR-959-964* has undergone relatively rapid evolutionary changes.

## Discussion

The spermatogenesis is a very complex and multistage process including cell proliferation, meiosis and differentiation and the spatial and temporal regulation of gene expression is of vital importance. Several recent reports have shown, that mammalian testes are characterized by a specific and unique set of miRNAs that can contribute to gene regulation [16–18, 21, 59]. Here we found that testes of *Drosophila melanogaster* are also characterized by the expression of 22 testis-specific miRNAs (Fig. 1A,B,C). Certainty, this set of miRNAs can be incomplete since we applied the quite strong requirement of at least 4-fold enrichment of miRNA abundance in testes in comparison to any other tissue. Indeed, some miRNAs in *Drosophila* (e.g. let-7, miRNA-34) are expressed at a relatively high level in several tissues and organs, and are reported to have specific functions in testes [24, 27, 29].

Intriguingly, 16 of the identified testis-specific miRNAs belong to only three miRNA clusters, which indicate the possibility of involvement of miRNAs in each cluster in co-regulation of pathways. We have shown, that the deletion of one of them, miR-959-964 cluster, is accompanied by improper spermatid individualization and male infertility (Fig. S2B; Fig. 2). Specifically, the Ics of mutant testes are moved in asynchronic and independent of each other manner downward the tails of the encysted bundle of spermatids (Fig. 2B,C; Fig. S3; Fig. S4). The spermatid nuclei in the *ΔmiR-959-964* testes are become to be separated before the replacement of histones to the protamine-like proteins in the elongated spermatids (Fig. 2D), that seemed to be the direct reason of the following failure of the individualization process. The expression of miR-959-964 cluster on the late stages of spermatogenesis is also indirectly confirmed by the fact that we failed to rescue the male sterility by the using of rescue transgene and several early-expressed GAL4 drivers.

To elucidate a possible mechanism of the observed phenotypic disorder, we identified the targets of miR-959-964 using the microarray technique followed by dual-luciferase assay validation (Figs 3, 4). One of the identified targets, *didum* has the largest number of miRNA recognitions sites (six sites for four miRNA, Table 1) and so it can be considered as the shared target of miR-959-964 cluster. It encodes the non-canonical, one-chain myosin of class V (MyoV) that is associated with the sperm nuclei during the maturation of the actin-rich ICs [60]. It has been suggested that MyoV contributes to the formation of the ICs and acts to coordinate as well as anchor these structures and other IC components. In *didum* mutant males, during the later stages of spermatogenesis, the ICs are poorly assembled and no mature sperms are produced [60]. It was also observed, that MyoV is expressed in the somatic cells of cyst and involved in maintaining of the germ cell differentiation [61]. Our data indicate that MyoV may also contribute to anchoring of nuclei of differentiating post-meiotic spermatids in proper positions near each other. Another interesting possibility is that miR-959-964 regulates the start point of coordinated initiation of the individualization of spermatids through the spatial and temporal control of *didum* expression. In the case of miR-959-964 deletion, the translation of *didum* may be launched too early that leads to the improper assembly of the ICs and failure of spermatid differentiation.

The levels of the expression of the other targets of miR-959-964, *for* and CG10512 encoding cAMP-dependent protein kinase and oxidoreductase respectively, are moderate in testis in comparison to other tissues. The functions of CG10512 are unknown while *for* is involved in feeding behavior of larvae and adult flies as well as short-term memory formation [62, 63]. It is not clear what testis-specific functions these target genes may have, and this question requires the future study. Alternative explanation of our observation based on the recent finding, that miR-959-964 in the head is involved in the circadian control of feeding [49]. If *for* is one of the miR-959-964 target that is responsible for such control, then our data may simply reflect the existence of residual and biologically unused regulation in adult testes.

It is considered that a high rate of evolution of testis-specific protein-coding genes is important for the speciation by reproductive isolation [64–66]. Recent reports indicate that several testis-specific miRNAs of mammals and flies are also characterized by rapid evolutionary changes [57, 58, 67] raising the possibility of their roles in reproductive isolation. In agreement with these fundings, we have shown that cluster of testis-specific miR-959-964 also undergoes the relatively rapid evolutionary process (Fig. 4). The species-specific differences of miRNA sequences and composition of the cluster have to result in the difference in the set of their targets and other proteins from the pathways in which the targets are involved. One can also speculate that distinctions of the protein set carried by mature sperm cells can prevent their ability to recognize and fertilize the eggs of other species. Although currently we are unable to directly verify this proposition, we noticed that one of the miR-959-964 targets is the gene *fdl* (Table 1), which is conserved among *Drosophilidae* species [68]. *fdl* encodes β-*N*-acetylhexosaminidase associated with the cytoplasmic membrane of mature sperms and it is also considered to be the putative receptor for glycoconjugates on the egg surface [69]. The distinction in the miR-959-964 structure can lead to the species-specific differences in the amount of Fdl receptor on the sperm membrane that may be one of the possible mechanisms in the reproductive isolation of species. If so, it is possible to speculate that a barrier to interspecies crosses may arises because sperms with low level *fdl* expression of one species unable to efficiently recognize the oocyte envelope of another species that was evolutionary fitted to be recognized by sperms with a high Fdl content.

Analysis of GO terms enrichment among functions of the down-regulated genes in *ΔmiR-959-964* testes revealed that many of them are involved in microtubule-based movement and the spermatid differentiation processes (Table S2; Table S3). There are two alternative explanations of this observation: either the down-regulation of such genes was caused by the failure in the spermatid differentiation process or the improper differentiation itself at least partially has occurred due to the genes down-regulation. Although it is hard to discriminate between these possibilities, the last alternative implies that genes down-regulation might be caused by the up-regulated miR-959-964 targets. Since we failed to find any common motifs for known transcriptional factors in the promoters of the down-regulated genes (data not shown), we suggest that their putative regulation by the up-regulated targets possibly occurred on the post-transcriptional level. But regardless of which alternative is true obviously that miR-959-964 is involved in direct and indirect expression regulation of the set of genes necessary for the differentiation of spermatid.

Are all miRNAs from miR-959-964 cluster contribute equally to the regulation of the spermatogenesis? In the recent genome-wide systematic study of effects of miRNA genes’ deletion on the fly phenotype was shown that the deletion of only miRNA-963 and miRNA-964 results to the male sterility while the deletion of miRNA-959, −960, −961 and −962 does not influence male fertility [56]. A little earlier the same group also reported that males carrying the deletion of the miR-959-962 part of the cluster are fertile like wild-type males [49]. Therefore, it is reasonable to conclude that miRNA-963 and miRNA-964 are major regulators of spermatogenesis. The indirect evidence of this conclusion come from our own observation that the insertion of CG31646^MI03191^ transgene occurred within *miRNA-963* resulted to the reduced fertility of young males and that this fertility partially restored with age (Fig. S2D, compare the 0-1 day old males with 2-4 day old males). Possibly, the MI03191 insertion disturbs the transcription and/or expression of only miR-963 and downstream miR-964, while the expression of the miRNAs located upstream of transgene insertion site are affected much lesser. In this way, the partial restoration of male fertility with age may occurred due to the decelerated rate of miRNA-959, −960, −961 and −963 accumulation and total absence of miRNA-963 and miR-964.

Because of transcriptional blockage, at the late stage of spermatogenesis the proteins are produced through the translation of mRNA synthesized only at the previous stage and regulation of gene expression occurred predominantly at the post-transcriptional level [29, 30]. Although there are several mechanisms of post-transcriptional gene expression regulation during late spermatogenesis are known [29, 30], it is reasonable to propose that miRNAs can also contribute to this process. To the best of our knowledge, the only example of the known involvement of miRNAs in regulation at the late stages of the spermatogenesis is the repression of transition protein gene *Tnp2* by miRNA-122 in mouse [21]. Although the precise mechanism of miR-959-964 involvement in spermatogenesis still remains to be investigated, here we report that it may be involved in the control of the spermatid individualization process in late spermatogenesis in *D. melanogaster*. *Didum*, one of the identified targets of miR-959-964, is the most plausible candidate for being a gene, that is responsible for spermatid individualization. 3’-UTR mRNA of *didum* carries 6 miRNA recognition sites for miR-959-964 including 1 site for miRNA-963 and 2 sites for miRNA-964 (Table 1). This rises the possibility that miR-959-964 may control the late spermatogenesis by regulating *didum* expression by miRNA-963/miRNA-964 that require the future investigation.

## Acknowledgements

We wish to thank R. Renkawitz-Pohl and C. Rathke for providing the anti-histones antibodies and *ProtB-eGFP* transgenic fly strain, J. Bischof for providing pUASTattB vector, Yu. Abramov for assistance with genetic crosses for the FLP-FRT mediated deletion, and Center of Biologically Active Compounds (VIGG RAS, Moscow, Russia) for microarray hybridization. We thank V. Gvozdev and A. Stolyarenko for the useful discussion and proof-editing of the manuscript. The research was supported by Russian Foundation for Basic Research (12-04-31352).

## References

1. Altuvia Y, Landgraf P, Lithwick G, Elefant N, Pfeffer S, Aravin A, Brownstein MJ, Tuschl T, Margalit H: Clustering and conservation patterns of human microRNAs. Nucleic Acids Res 2005, 33:2697–706.

2. Ryazansky SS, Gvozdev VA, Berezikov E: Evidence for post-transcriptional regulation of clustered microRNAs in Drosophila. BMC Genomics 2011, 12(1471-2164 (Electronic)):371.

3. Merchan F, Boualem A, Crespi M, Frugier F: Plant polycistronic precursors containing non-homologous microRNAs target transcripts encoding functionally related proteins. Genome Biol 2009, 10:R136.

4. Bartel DP: MicroRNAs: genomics, biogenesis, mechanism, and function. Cell 2004, 116:281– 97.

5. Filipowicz W, Bhattacharyya SN, Sonenberg N: Mechanisms of post-transcriptional regulation by microRNAs: are the answers in sight? Nat Rev Genet 2008, 9:102–14.

6. Guo H, Ingolia NT, Weissman JS, Bartel DP: Mammalian microRNAs predominantly act to decrease target mRNA levels. Nature 2010, 466:835–40.

7. Hendrickson DG, Hogan DJ, McCullough HL, Myers JW, Herschlag D, Ferrell JE, Brown PO: Concordant regulation of translation and mRNA abundance for hundreds of targets of a human microRNA. PLoS Biol 2009, 7:e1000238.

8. Ryazansky SS, Mikhaleva EA, Olenkina O V: Essential Functions of microRNAs in Animal Reproductive Organs. Mol Biol 2014, 48:319–331.

9. Wang L, Xu C: Role of microRNAs in mammalian spermatogenesis and testicular germ cell tumors. Reproduction 2015, 149:R127–R137.

10. Kotaja N, Bhattacharyya SN, Jaskiewicz L, Kimmins S, Parvinen M, Filipowicz W, Sassone-Corsi P: The chromatoid body of male germ cells: similarity with processing bodies and presence of Dicer and microRNA pathway components. Proc Natl Acad Sci USA 2006, 103:2647–52.

11. Hayashi K, Chuva de Sousa Lopes SM, Kaneda M, Tang F, Hajkova P, Lao K, O’Carroll D, Das PP, Tarakhovsky A, Miska EA, Surani MA: MicroRNA biogenesis is required for mouse primordial germ cell development and spermatogenesis. PLoS One 2008, 3:e1738.

12. Korhonen HM, Meikar O, Yadav RP, Papaioannou MD, Romero Y, Da Ros M, Herrera PL, Toppari J, Nef S, Kotaja N: Dicer is required for haploid male germ cell differentiation in mice. PLoS One 2011, 6:e24821.

13. Maatouk DM, Loveland KL, McManus MT, Moore K, Harfe BD: Dicer1 is required for differentiation of the mouse male germline. Biol Reprod 2008, 79:696–703.

14. Romero Y, Meikar O, Papaioannou MD, Conne B, Grey C, Weier M, Pralong F, De Massy B, Kaessmann H, Vassalli J-D, Kotaja N, Nef S: Dicer1 depletion in male germ cells leads to infertility due to cumulative meiotic and spermiogenic defects. PLoS One 2011, 6:e25241.

15. Wu Q, Song R, Ortogero N, Zheng H, Evanoff R, Small CL, Griswold MD, Namekawa SH, Royo H, Turner JM, Yan W: The RNase III enzyme DROSHA is essential for microRNA production and spermatogenesis. J Biol Chem 2012, 287:25173–90.

16. Linsen SE V, de Wit E, de Bruijn E, Cuppen E: Small RNA expression and strain specificity in the rat. BMC Genomics 2010, 11(1471-2164 (Electronic)):249.

17. Marcon E, Babak T, Chua G, Hughes T, Moens PB: miRNA and piRNA localization in the male mammalian meiotic nucleus. Chromosome Res 2008, 16:243–60.

18. McIver SC, Stanger SJ, Santarelli DM, Roman SD, Nixon B, McLaughlin EA: A unique combination of male germ cell miRNAs coordinates gonocyte differentiation. PLoS One 2012, 7:e35553.

19. Ro S, Park C, Sanders KM, McCarrey JR, Yan W: Cloning and expression profiling of testis-expressed microRNAs. Dev Biol 2007, 311:592–602.

20. Yan N, Lu Y, Sun H, Qiu W, Tao D, Liu Y, Chen H, Yang Y, Zhang S, Li X, Ma Y: Microarray profiling of microRNAs expressed in testis tissues of developing primates. J Assist Reprod Genet 2009, 26:179–86.

21. Yu Z, Raabe T, Hecht NB: MicroRNA Mirn122a reduces expression of the posttranscriptionally regulated germ cell transition protein 2 (Tnp2) messenger RNA (mRNA) by mRNA cleavage. Biol Reprod 2005, 73:427–33.

22. Novotny GW, Sonne SB, Nielsen JE, Jonstrup SP, Hansen M a, Skakkebaek NE, Rajpert-De Meyts E, Kjems J, Leffers H: Translational repression of E2F1 mRNA in carcinoma in situ and normal testis correlates with expression of the miR-17-92 cluster. Cell Death Differ 2007, 14:879–82.

23. Niu Z, Goodyear SM, Rao S, Wu X, Tobias JW, Avarbock MR, Brinster RL: MicroRNA-21 regulates the self-renewal of mouse spermatogonial stem cells. Proc Natl Acad Sci USA 2011, 108:12740–5.

24. Toledano H, D’Alterio C, Czech B, Levine E, Jones DL: The let-7-Imp axis regulates ageing of the Drosophila testis stem-cell niche. Nature 2012, 485:605–10.

25. Bao J, Li D, Wang L, Wu J, Hu Y, Wang Z, Chen Y, Cao X, Jiang C, Yan W, Xu C: MicroRNA-449 and microRNA-34b/c function redundantly in murine testes by targeting E2F transcription factor-retinoblastoma protein (E2F-pRb) pathway. J Biol Chem 2012, 287:21686–98.

26. Bouhallier F, Allioli N, Lavial F, Chalmel F, Perrard M-H, Durand P, Samarut J, Pain B, Rouault J-P: Role of miR-34c microRNA in the late steps of spermatogenesis. RNA 2010, 16:720–31.

27. Liang X, Zhou D, Wei C, Luo H, Liu J, Fu R, Cui S: MicroRNA-34c enhances murine male germ cell apoptosis through targeting ATF1. PLoS One 2012, 7:e33861.

28. Liu W-M, Pang RTK, Chiu PCN, Wong BPC, Lao K, Lee K-F, Yeung WSB: Sperm-borne microRNA-34c is required for the first cleavage division in mouse. Proc Natl Acad Sci USA 2012, 109:490–4.

29. Idler RK, Yan W: Control of messenger RNA fate by RNA-binding proteins: an emphasis on mammalian spermatogenesis. J Androl 2012, 33:309–37.

30. Kleene KC: Patterns, mechanisms, and functions of translation regulation in mammalian spermatogenic cells. Cytogenet Genome Res 2003, 103:217–24.

31. Langmead B, Trapnell C, Pop M, Salzberg SL: Ultrafast and memory-efficient alignment of short DNA sequences to the human genome. Genome Biol 2009, 10:R25.

32. Meyer LR, Zweig AS, Hinrichs AS, Karolchik D, Kuhn RM, Wong M, Sloan CA, Rosenbloom KR, Roe G, Rhead B, Raney BJ, Pohl A, Malladi VS, Li CH, Lee BT, Learned K, Kirkup V, Hsu F, Heitner S, Harte RA, Haeussler M, Guruvadoo L, Goldman M, Giardine BM, Fujita PA, Dreszer TR, Diekhans M, Cline MS, Clawson H, Barber GP, et al.: The UCSC Genome Browser database: extensions and updates 2013. Nucleic Acids Res 2013, 41(Database issue):D64–9.

33. Kozomara A, Griffiths-Jones S: miRBase: integrating microRNA annotation and deep-sequencing data. Nucleic Acids Res 2011, 39(Database issue):D152–7.

34. Parks AL, Cook KR, Belvin M, Dompe N a, Fawcett R, Huppert K, Tan LR, Winter CG, Bogart KP, Deal JE, Deal-Herr ME, Grant D, Marcinko M, Miyazaki WY, Robertson S, Shaw KJ, Tabios M, Vysotskaia V, Zhao L, Andrade RS, Edgar K a, Howie E, Killpack K, Milash B, Norton A, Thao D, Whittaker K, Winner M a, Friedman L, Margolis J, et al.: Systematic generation of high-resolution deletion coverage of the Drosophila melanogaster genome. Nat Genet 2004, 36:288– 92.

35. Dietzl G, Chen D, Schnorrer F, Su K-C, Barinova Y, Fellner M, Gasser B, Kinsey K, Oppel S, Scheiblauer S, Couto A, Marra V, Keleman K, Dickson BJ: A genome-wide transgenic RNAi library for conditional gene inactivation in Drosophila. Nature 2007, 448:151–6.

36. Jayaramaiah Raja S, Renkawitz-Pohl R: Replacement by Drosophila melanogaster protamines and Mst77F of histones during chromatin condensation in late spermatids and role of sesame in the removal of these proteins from the male pronucleus. Mol Cell Biol 2005, 25:6165–77.

37. Kibanov M V, Egorova KS, Ryazansky SS, Sokolova OA, Kotov AA, Olenkina OM, Stolyarenko AD, Gvozdev VA, Olenina L V: A novel organelle, the piNG-body, in the nuage of Drosophila male germ cells is associated with piRNA-mediated gene silencing. Mol Biol Cell 2011, 22:3410–9.

38. White-Cooper H: Spermatogenesis: analysis of meiosis and morphogenesis. Methods Mol Biol 2004, 247(1064-3745 (Print)):45–75.

39. Smyth GK: Limma: linear models fro microarray data. In Bioinformatics and Computational Biology Solutions using R and Bioconductor. edited by Gentleman R, Carey V, Dudoit S, Irizarry R, Huber W. New York: Springer; 2005:397–420.

40. Lyne R, Smith R, Rutherford K, Wakeling M, Varley A, Guillier F, Janssens H, Ji W, Mclaren P, North P, Rana D, Riley T, Sullivan J, Watkins X, Woodbridge M, Lilley K, Russell S, Ashburner M, Mizuguchi K, Micklem G: FlyMine: an integrated database for Drosophila and Anopheles genomics. Genome Biol 2007, 8:R129.

41. Bischof J, Maeda RK, Hediger M, Karch F, Basler K: An optimized transgenesis system for Drosophila using germ-line-specific phiC31 integrases. Proc Natl Acad Sci USA 2007, 104:3312–7.

42. Karolchik D, Hinrichs AS, Furey TS, Roskin KM, Sugnet CW, Haussler D, Kent WJ: The UCSC Table Browser data retrieval tool. Nucleic Acids Res 2004, 32(Database issue):D493–6.

43. Felsenstein J: PHYLIP (Phylogeny Inference Package) version 3.6. 2005.

44. Hofacker IL, Fontana W, Stadler PF, Bonhoeffer LS, Tacker M, Schuster P: Fast folding and comparison of RNA secondary structures. Monatshefte fr Chemie Chem Mon 1994, 125:167– 188.

45. Bonnet E, Wuyts J, Rouzé P, Van de Peer Y: Evidence that microRNA precursors, unlike other non-coding RNAs, have lower folding free energies than random sequences. Bioinformatics 2004, 20:2911–7.

46. Kimura M: A simple method for estimating evolutionary rates of base substitutions through comparative studies of nucleotide sequences. J Mol Evol 1980, 16:111–20.

47. Stajich JE, Block D, Boulez K, Brenner SE, Chervitz SA, Dagdigian C, Fuellen G, Gilbert JGR, Korf I, Lapp H, Lehväslaiho H, Matsalla C, Mungall CJ, Osborne BI, Pocock MR, Schattner P, Senger M, Stein LD, Stupka E, Wilkinson MD, Birney E: The Bioperl toolkit: Perl modules for the life sciences. Genome Res 2002, 12:1611–8.

48. Kinoshita T, Hanazawa T, Nohata N, Okamoto Y, Seki N: The functional significance of microRNA-375 in human squamous cell carcinoma: aberrant expression and effects on cancer pathways. J Hum Genet 2012, 57:556–63.

49. Vodala S, Pescatore S, Rodriguez J, Buescher M, Chen Y-W, Weng R, Cohen SM, Rosbash M: The oscillating miRNA 959-964 cluster impacts Drosophila feeding time and other circadian outputs. Cell Metab 2012, 16:601–12.

50. Celniker SE, Dillon LAL, Gerstein MB, Gunsalus KC, Henikoff S, Karpen GH, Kellis M, Lai EC, Lieb JD, MacAlpine DM, Micklem G, Piano F, Snyder M, Stein L, White KP, Waterston RH: Unlocking the secrets of the genome. Nature 2009, 459:927–30.

51. Fabian L, Brill JA: Drosophila spermiogenesis: Big things come from little packages. Spermatogenesis 2012, 2:197–212.

52. Fabrizio JJ, Hime G, Lemmon SK, Bazinet C: Genetic dissection of sperm individualization in Drosophila melanogaster. Development 1998, 125:1833–43.

53. Jayaramaiah Raja S, Renkawitz-Pohl R: Replacement by Drosophila melanogaster protamines and Mst77F of histones during chromatin condensation in late spermatids and role of sesame in the removal of these proteins from the male pronucleus. Mol Cell Biol 2005, 25:6165–77.

54. Ruby JG, Stark A, Johnston WK, Kellis M, Bartel DP, Lai EC: Evolution, biogenesis, expression, and target predictions of a substantially expanded set of Drosophila microRNAs. Genome Res 2007, 17:1850–64.

55. Szuplewski S, Kugler J-M, Lim SF, Verma P, Chen Y-W, Cohen SM: MicroRNA transgene overexpression complements deficiency-based modifier screens in Drosophila. Genetics 2012, 190:617–26.

56. Chen Y-W, Song S, Weng R, Verma P, Kugler J-M, Buescher M, Rouam S, Cohen SM: Systematic Study of Drosophila MicroRNA Functions Using a Collection of Targeted Knockout Mutations. Dev Cell 2014, 31:784–800.

57. Guo X, Su B, Zhou Z, Sha J: Rapid evolution of mammalian X-linked testis microRNAs. BMC Genomics 2009, 10(1471-2164 (Electronic)):97.

58. Zhang R, Peng Y, Wang W, Su B: Rapid evolution of an X-linked microRNA cluster in primates. Genome Res 2007, 17:612–7.

59. Yan N, Lu Y, Sun H, Tao D, Zhang S, Liu W, Ma Y: A microarray for microRNA profiling in mouse testis tissues. Reproduction 2007, 134:73–9.

60. Mermall V, Bonafé N, Jones L, Sellers JR, Cooley L, Mooseker MS: Drosophila myosin V is required for larval development and spermatid individualization. Dev Biol 2005, 286:238–55.

61. Joti P, Ghosh-Roy A, Ray K: Dynein light chain 1 functions in somatic cyst cells regulate spermatogonial divisions in Drosophila. Sci Rep 2011, 1(2045-2322 (Electronic)):173.

62. Pereira HS, Sokolowski MB: Mutations in the larval foraging gene affect adult locomotory behavior after feeding in Drosophila melanogaster. Proc Natl Acad Sci USA 1993, 90:5044–6.

63. Kent CF, Daskalchuk T, Cook L, Sokolowski MB, Greenspan RJ: The Drosophila foraging gene mediates adult plasticity and gene-environment interactions in behaviour, metabolites, and gene expression in response to food deprivation. PLoS Genet 2009, 5:e1000609.

64. Turner LM, Chuong EB, Hoekstra HE: Comparative analysis of testis protein evolution in rodents. Genetics 2008, 179:2075–89.

65. Swanson WJ, Vacquier VD: The rapid evolution of reproductive proteins. Nat Rev Genet 2002, 3:137–44.

66. Li VC, Davis JC, Lenkov K, Bolival B, Fuller MT, Petrov DA: Molecular evolution of the testis TAFs of Drosophila. Mol Biol Evol 2009, 26:1103–16.

67. Mohammed J, Bortolamiol-Becet D, Flynt AS, Gronau I, Siepel A, Lai EC: Adaptive evolution of testis-specific, recently evolved, clustered miRNAs in Drosophila. RNA 2014.

68. Intra J, Cenni F, Pavesi G, Pasini M, Perotti M-E: Interspecific analysis of the glycosidases of the sperm plasma membrane in Drosophila. Mol Reprod Dev 2009, 76:85–100.

69. Cattaneo F, Ogiso M, Hoshi M, Perotti ME, Pasini ME: Purification and characterization of the plasma membrane glycosidases of Drosophila melanogaster spermatozoa. Insect Biochem Mol Biol 2002, 32:929–41.

